# Algebraic principle of natural bases from the standard genetic code degeneracy

**DOI:** 10.1101/2023.04.10.536215

**Authors:** Yang Yu

## Abstract

The symmetry and group in degeneracy of the standard genetic code (SGC) have been studied. However, the core role of equations of degree n with one unknown between symmetry and group theory has been ignored. In this study, algebraic concept was employed to abstract all genetic codons in the SGC table into equations of degree n with one unknown, and the bases in the third position of nucleotide triplets are these equations solutions. Basing upon the analysis of natural bases permutation groups and natural bases as the unit roots, some results were found that the characteristics of solvable groups imply the amount of natural bases, the relation between the first two bases in nucleotide triplets is the algebraic multiplication binary operation, and natural bases have the significance of complex number. These results gave natural bases the significance of number in the complex plane, and clarified the amount and operation in the mathematical sense, which would contribute to understand the origin and evolution of genetic code.

## Introduction

The degeneracy rules of the standard genetic code (SGC) is the existence of silent or synonymous mutations (Hunt RC et al., 2014; Lehmann J et al., 2008; Ye S et al., 2022). The specificity of amino acid is determined by the first two bases in nucleotide triplets, while the variability of the third base implies the substitutions of four natural bases uracil (U), cytosine (C), adenine (A) and guanine (G) in messenger RNA (mRNA). The symmetry and group theory in the SGC degeneracy have been studied (Antillon A et al., 1985; Bashford JD et al., 1998; Fimmel E et al., 2018; Hornos JE et al., 1993; Jestin JL, 2010; Rietman EA et al., 2011; Sanchez R et al., 2005; Sanchez R et al., 2006). The focus of those studies is the representations of the SGC and generalized genetic code (GGC), whose purpose is to find the digital representation or model of genetic information. Nevertheless, the core role of equations of degree n with one unknown between symmetry and group theory has been ignored in the past research all the time.

In mathematics, the characteristics of unchanged output with changed input are symmetry. Symmetry could be studied by group theory. Group is one kind of algebraic structure basing upon the set and operation, and viewed as one of the basic structures in mathematics. In fact, from the beginning of groundbreaking theory by mathematician Galois, the solutions of equation of degree n with one unknown, symmetry and permutation group have been integrated together tightly. In general, permutation group is a subgroup of the symmetric group. Group theory is from the research of solutions of equation of degree n with one unknown, and solutions permutations of these equations imply symmetry and group theory. So equations and their solutions play a central role in the understanding of symmetry and group (Emil Artin et al., 2012; Harold M. Edwards, 1984; Ian Stewart, 2022; John M. Howie, 2007).

Degeneracy of the SGC includes double, triple and quadruple degeneracy. In these degeneracies, synonymous mutations mean only base substitutions but no amino acid changes, which determine a certain of symmetry. These base substitutions constitute the permutations and symmetry in the mathematical sense, just like the permutations and symmetry of equation solutions. For example of the amino acid Asp, GAU and GAC include the substitutions between U and C, which do not change amino acid. So, GAU and GAC determine the symmetry about Asp, and U and C establish the permutations in the third base of nucleotide triplet.

Here, basing upon the equations of degree n with one unknown and their solutions, mathematical concept of set, permutation, symmetric group, group structure, solvable group, binary operation and unit root was employed to elaborate the algebraic principle of natural bases for the amount, multiplication operation and complex number significance from the SGC degeneracy.

## Results and Discussion

### Sets from bases substitutions and permutations in symmetric group

In mathematics, symmetry and permutation group from equations of degree n with one unknown are established on the basis of bijection on set itself, which is the mathematical foundation of the subsequent analysis.

In double degeneracy of the SGC, there are the substitutions between purines or pyrimidines, for example, GAU and GAC determine Asp while GAA and GAG determine Glu. In triple degeneracy, there are the substitutions among U, C and A, for only one example, AUU, AUC and AUA determine Ile. In quadruple degeneracy, there are the substitutions among U, C, A and G, for example, GUU, GUC, GUA and GUG determine Val. Consequently, there are sets from the bases substitutions in the third position of the genetic codons: {U,C} for Asp, {A,G} for Glu, {U,C,A} for Ile and {U,C,A,G} for Val.

For the amino acid of Asp, U and C determine one set {U,C} which includes two elements. In the {U,C} set, considering a collection group of all permutations from bijections of {U,C} set into itself, the group is the symmetric group which could be notated as S_n_ (n=the number of elements in a given set). In the {U,C} set, there are two permutations (the number of permutations is n factorial, n=the number of element in a given set, 2!=2*1=2), {U to U, C to C} and {U to C, C to U}. The permutation of {U to U, C to C} is identity (notated as e in group theory) which is no substitutions. The permutation of {U to C, C to U} is exchange or transposition permutation between U and C, which is synonymous mutation in genetic codon, notated as (U C) according to group theory. According to the corresponding bases substitutions and permutation group theory, similar results could be obtained in the set {A,G} for Glu, {U,C,A} for Ile and {U,C,A,G} for Val (the number of permutations are 2!=2*1=2, 3!=3*2*1=6 and 4!=4*3*2*1=24 respectively).

### Mathematical equations and group structures in the SGC table

Considering the knowledge of equation solutions permutations and group theory, GAU and GAC coding Asp could be abstracted into a mathematical equation of GAx^2^=Asp. U and C are two solutions of equation GAx^2^=Asp: x_1_=U and x_2_=C. The two permutations of e and (U C) establish the symmetric group S_2_. The only one normal subgroup of S_2_ is e. Meanwhile, S_2_ is also a cyclic group C_2_ (the cyclic group of order 2). The permutation of (U C) is the generator of C_2_, that is (U C)(U C) or (U C)^2^=e. So, the group structure is S_2_⊃e, and the normal subgroup chain is S_2_ ▷ e. Obviously, the {U,C} group from GAx^2^=Asp is isomorphic to the {A,G} group from GAx^2^=Glu.

Similarly, AUU, AUC and AUA coding Ile could be abstracted into a equation of AUx^3^=Ile. U, C and A are three solutions of equation AUx^3^=Ile: x_1_=U, x_2_=C, x_3_=A. The group structure is S_3_⊃A_3_⊃e, and the normal subgroup chain is S_3_ ▷ A_3_ ▷ e. A_3_ is an alternating group which is the group of all even permutations of a finite set. That is, all even permutations of {U,C,A} establish the group A_3_. Coincidentally, A_3_ is also a cyclic group C_3_. C_3_ includes three permutations of e, (U C A) and (U A C). (U C A) means {U to C, C to A, A to U} while (U A C) means {U to A, A to C, C to U}. (U C A)^2^=(U A C), and (U C A)^3^=e, so (U C A) is one generator of C_3_. Meanwhile, (U A C)^2^=(U C A), (U A C)^3^=e, so (U A C) is the other generator of C_3_. And also, GUU, GUC, GUA and GUG coding Val could be abstracted into a mathematical equation of GUx^4^=Val. U, C, A and G are four solutions of equation GUx^4^=Val: x_1_=U, x_2_=C, x_3_=A, x_4_=G. The group structure is S_4_⊃A_4_⊃K_4_⊃{e, (U C)(A G)}⊃e, and the normal subgroup chain is S_4_ ▷A_4_ ▷K_4_ ▷ e. K_4_ is the Klein four-group, and it’s permutations include {e, (U C)(A G), (U A)(C G), (U G)(C A)}.

### The characteristics of solvable groups imply the amount of natural bases

Above analysis could cover different situations of mathematical equations and group structures in the degeneracy rules from the SGC table. Consequently, all genetic codons in the SGC table could be abstracted into a series of equations of degree n with one unknown in Table 1. There are five kinds of solutions organized by 4 natural bases for the 20 natural amino acids: G, U/C, A/G, U/C/A and U/C/A/G. There are no degeneracy in AUG=Met and UGG=Trp. Degeneracy includes four kinds of the solutions organization with U/C, A/G, U/C/A and U/C/A/G, and U/C, A/G and U/C/A/G organizations cover the absolute majority in degeneracy of the SGC (U/C/A is special only for the amino acid Ile). Significantly, in the above symmetric groups, S_2_, S_3_ and S_4_ are very special, and they are solvable groups. When n≥5, S_n_ are the unsolvable groups, so there are no regular algebraic solutions for equations of degree n (n≥5) with one unknown (Harold M. Edwards, 1984). The amount of natural bases for the natural 20 amino acids is 4, and this amount could be implied by the characteristics of solvable groups. The amount of 4 not only maximizes the efficiency of the SGC (compared with n<4) but also ensures the third position of all genetic codons in the SGC table solvable (compared with n>4).

**Table 1.**
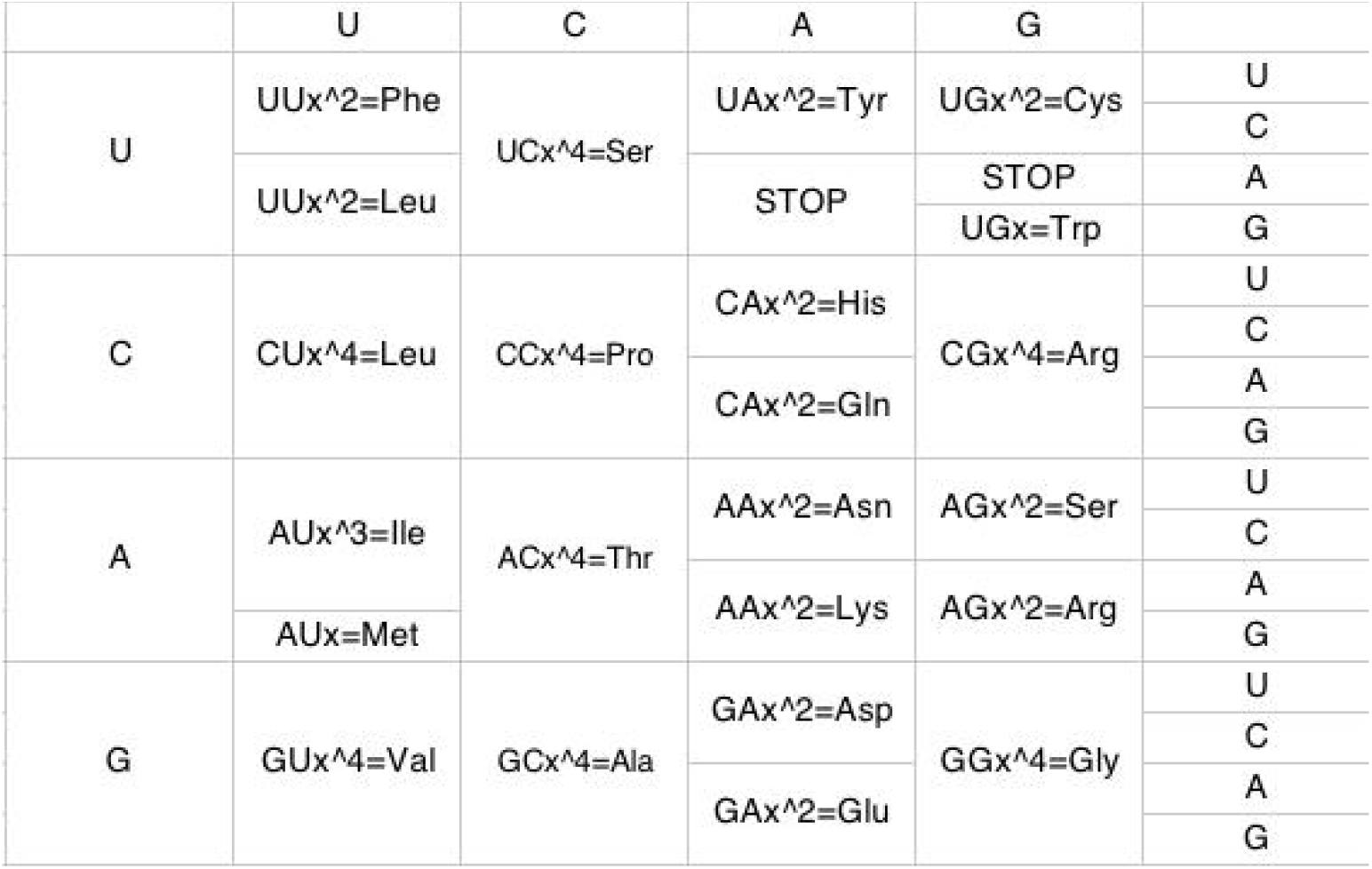
All genetic codons as equations of degree n with one unknown in the SGC table.

### The multiplication operation between the first two bases of nucleotide triplets

The splitting field and field extension in the solvable equations must be paid attention. In fact, if there are any solutions for the equation of degree n with one unknown, field extension would exist, which is the Galois extension (Emil Artin et al., 2012; Harold M. Edwards, 1984; Ian Stewart, 2022; John M. Howie, 2007). The solutions of GAx^2^=Glu, GAx^2^=Asp, AUx^3^=Ile and GUx^4^=Val are not from the original field of equation coefficients themselves but from the splitting field and field extension which is the normal extension with the existence of normal subgroup. The groups of S_2_, S_3_ and S_4_ are solvable, so the corresponding field extensions are effective. In the equations from the SGC table, the first two bases in nucleotide triplets could be looked as the equation coefficients. There are 16 kinds of equation coefficients, that is UU, UC, UA, UG, CU, CC, CA, CG, AU, AC, AA, AG, GU, GC, GA and GG, which are all dinucleotides. Nevertheless, all equations solutions are mononucleotides, including U, C, A and G. There is a significant difference between the original field from equations coefficients and the splitting field from equations solutions after field extension: dinucleotide in the first two bases of nucleotide triplets while mononucleotide in the third position. There is no solutions in the original field from equation coefficients, and only mononucleotides could be the solutions of equations. There is the process of rooting from the original field to the splitting field in the equations of degree n with one unknown. Inspired by this field analysis, the viewpoint could be prompted that the relation between the first two bases in nucleotide triplets is the algebraic multiplication binary operation.

### Natural bases as the unit roots of cyclotomic equations

Considering 4 bases and 20 amino acids are natural in our world, they could be regarded as the unit bases and amino acids. Consequently, all mathematical equations from the SGC table, such as GAx^2^=Glu, GAx^2^=Asp, AUx^3^=Ile and GUx^4^=Val patterns, could be considered as cyclotomic equation x^n^=1 (n=2, 3 or 4, and stands for the double, triple or quadruple degeneracy respectively) in mathematics. In fact, this perspective is reasonable because of fundamentality of 4 bases and 20 amino acids in natural world, just like elementary and abstract meaning of the number 1 in mathematics. The natural bases and amino acids are not only the practical things but also abstract unit of homogeneity in the algebraic abstract sense. All solutions of cyclotomic equation distribute on the n-bisector of the unit circle on the complex plane (argand plane), and these solutions are unit roots. In this analysis, the order of the roots are followed according to the SGC table: U, C, A and G. In x^n^=1, x is unit root and x=cos(2kπ/n)+i sin(2kπ/n), k=1, 2, …, n-1, n. When k and n are relatively prime, root x is the primitive unit root which is also the generator of cyclic group from multiplication of unit root. Moreover, there must be a unit root of 1 in any cyclotomic equation, which is the e of the cyclic group. In fact, the base G is very special for the evolution of the SGC, because G is the only solution in initiation codon AUx=Met, which is no existence of degeneracy but determinacy. So is the x=G in UGx=Trp. That is, when n=1, cyclotomic equation is x=1. So the distribution of G as the unit root on the unit circle overlaps with 1 (G=1) in AUx=Met and UGx=Trp (Figure 1a), which coincides with the distribution situations in GAx^2^=Glu and GUx^4^=Val in the following analysis.

**Figure 1.**
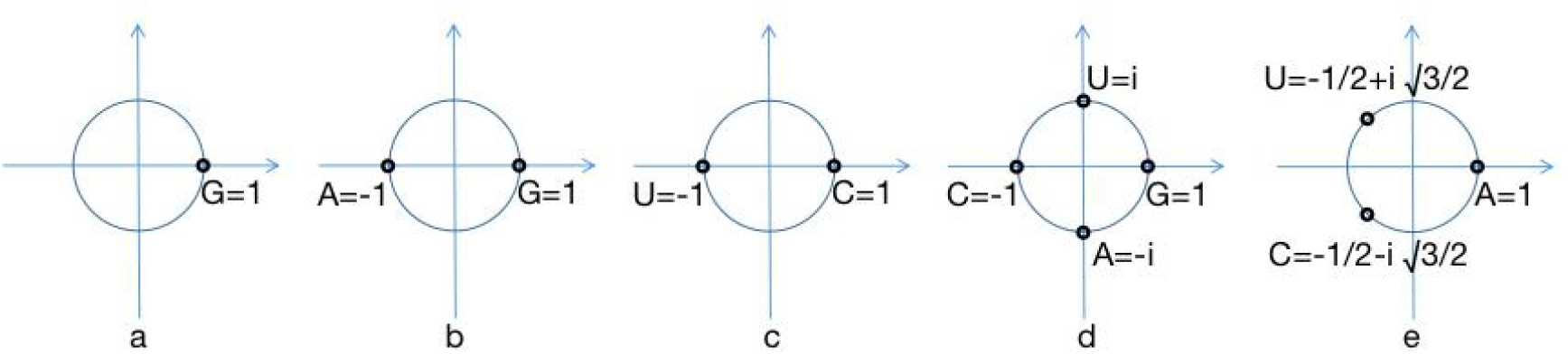
Natural bases as the unit roots of cyclotomic equation. a. x=1 such as AUx=Met pattern. b. x^2^=1 such as GAx^2^=Glu pattern. c. x^2^=1 such as GAx^2^=Asp pattern. d. x^4^=1 such as GUx^4^=Val pattern. e. x^3^=1 such as AUx^3^=Ile pattern.

x^2^=1 and x^4^=1 cover the absolute majority in degeneracy of the SGC. In the pattern of GAx^2^=Glu, x_1_=A and x_2_=G. When n=2 and k=1, k and n are relatively prime, and x_1_=cos π+i sinπ=-1. When k=2, k and n are not relatively prime, and x_2_=cos(2π)+i sin(2π)=1. So A=-1 is the primitive unit root, and cyclotomic polynomial is x+1. G=1. (Figure 1b) (−1)^2^=1, so A^2^=G=1=e. The group of {A, G} is a cyclic group, and the generator is A. Similarly, U and C are two roots of GAx^2^=Asp. When k=1, x_1_=cosπ+i sinπ=-1. When k=2, x_2_=cos(2π)+i sin(2π)=1. So U=-1 is the primitive unit root, and cyclotomic polynomial is x+1. C=1. (Figure 1c) U^2^=C=1=e. The group of {U, C} is a cyclic group, and the generator is U. Obviously, the {U, C} group is isomorphic to the {A, G} group. In the pattern of GUx^4^=Val, x_1_=U, x_2_=C, x_3_=A, x_4_=G, and x_1_=cos(π/2)+i sin(π/2)=i, x_2_=cosπ+i sinπ=-1, x_3_=cos(3π/2)+i sin(3π/2)=-i, x_4_=cos(2π)+i sin(2π)=1. So U=i, C=-1, A=-i, and G=1. (Figure 1d) U=i and A=-i are primitive unit roots, and cyclotomic polynomial is x^2^+1. U^2^=C, U^3^=A, U^4^=G=1=e, A^2^=C, A^3^=U and A^4^=G=1=e. The group of {U, C, A, G} is a cyclic group, and the two generators are U and A.

At last, x^3^=1 is for only one case of Ile, and is very special. In the pattern of AUx^3^=Ile, x_1_=U, x_2_=C and x_3_=A. When n=3 and k=1, k and n are relatively prime, x_1_=cos(2 π/3)+i sin(2π/3)=-1/2+i √3/2. When k=2, k and n are also relatively prime, x_2_=cos(4π/3)+i sin(4 π/3)=-1/2-i √3/2. When k=3, k and n are not relatively prime, x_3_=cos(2π)+i sin(2π)=1. So, U=-1/2+i √3/2, C=-1/2-i √3/2 and A=1. (Figure 1e) U and C are the primitive unit roots, and cyclotomic polynomial is x^2^+x+1. U^2^=C, U^3^=A=1=e, C^2^=U and C^3^=A=1=e. The group of {U, C, A} is a cyclic group, and the two generators are U and C.

### Complex number significance of natural bases

In mathematics, complex number is the largest set of number, in which many mathematical and physical problems could be solved effectively. Complex number includes two dimensions, real number part and imaginary number part, which establish the complex plane (Ian Stewart, 2022). Locating natural bases on the unit circle on the complex plane could provide the sight determining natural bases as complex number significance.

x^2^=1 and x^4^=1 cover the absolute majority in the SGC degeneracy. There are two kinds of unit roots for x^2^=1 from the SGC table: A and G (all purines), or U and C (all pyrimidines). Only two purines or pyrimidines appear in the unit roots, and no existence of one purine mixed with one pyrimidine. The {U, C} group and {A, G} group are relatively isomorphic. So there is only one dimension for the unit roots of cyclotomic equation x^2^=1, and these unit roots are set by the unit of real number 1 and -1. In x^2^=1, U or A is the primitive unit root. In the situations of x^4^=1, all two purines and two pyrimidines appear in the unit roots. Besides the unit of real number 1 and -1 as the first dimension, the unit of imaginary number i and -i, as the second dimension, are introduced to show the bases as the unit roots on unit circle. So there are two dimensions for U, C, A and G as the unit roots of cyclotomic equation x^4^=1. Significantly, the second dimension (i and -i) was determined by the primitive unit roots (U and A, matching each other with weak hydrogen bonds). In x^3^=1, U and C are all primitive unit roots which are set by complex number (mixed with real part and imaginary part).

This study is the process of algebraic abstraction for the SGC degeneracy to find the algebraic principle of natural bases from the SGC degeneracy in the aid of the equations of degree n with one unknown and their solutions. Basing upon the analyzed results of natural bases permutation groups and natural bases as the unit roots, I found that the characteristics of solvable groups imply the amount of natural bases, the relation between the first two bases in nucleotide triplets is the algebraic multiplication binary operation, and natural bases have the significance of complex number. These results gave natural bases the significance of number in the complex plane, and clarified the amount and operation in the mathematical sense, which would contribute to understand the origin and evolution of genetic code. Meanwhile, the methods of equations in this study capture the core and origin of the understanding of symmetry and group, which could be the inspiration for the future biological research by the mathematical method.

## Methods

Symmetry about no change of amino acid with substitutions of the third base in nucleotide triplets was clarified. Sets and permutations from these base substitutions were defined. Equation solutions and base permutations were linked, and symmetric group structures were analyzed. All genetic codons in the SGC table were abstracted into a series of equations of degree n with one unknown. Furthermore, according to the idea of abstract unit of natural 4 bases and 20 amino acids, the above mathematical equations were abstracted as cyclotomic equation. Natural bases as the unit roots of cyclotomic equation and corresponding algebraic property were analyzed.

## Acknowledgments

I would like to thank Jing Zhou for help and advice for comments on the manuscript.

## Competing interests

The author declare no competing interests.

## Notes

### Competing Interest Statement

The authors have declared no competing interest.

### Summary of Updates

Abstract and conclusion updated for text simplification and modification.

## References

Antillon A, Ortega-Blake I (1985) A group theory analysis of the ambiguities in the genetic code: on the existence of a generalized genetic code Journal of Theoretical Biology 112:757–769. https://doi.org/10.1016/s0022-5193(85)80059-x.

Bashford JD, Tsohantjis I, Jarvis PD (1998) A supersymmetric model for the evolution of the genetic code Proceeding of the National Academy of Sciences of the United States of America 95:987–992. https://doi.org/10.1073/pnas.95.3.987.

Emil Artin, Arthur N. Milgram (2012) Galois Theory Dover Publications 9780486158259.

Fimmel E, Strungmann L (2018) Mathematical fundamentals for the noise immunity of the genetic code Biosystems 164:186–198. https://doi.org/10.1016/j.biosystems.2017.09.007.

Harold M. Edwards (1984) Galois Theory Springer 9780387909806.

Hornos JE, Hornos YM (1993) Algebraic model for the evolution of the genetic code Physical Review Letters 71:4401–4404. https://doi.org/10.1103/PhysRevLett.71.4401.

Hunt RC, Simhadri VL, Iandoli M, Sauna ZE, Kimchi-Sarfaty C (2014) Exposing synonymous mutations Trends in Genetics 30:308–321. https://doi.org/10.1016/j.tig.2014.04.006.

Ian Stewart (2022) Galois Theory CRC Press 9781000644067.

Jestin JL (2010) A rationale for the symmetries by base substitutions of degeneracy in the genetic code Biosystems 99:1–5. https://doi.org/10.1016/j.biosystems.2009.07.009.

John M. Howie (2007) Fields and Galois Theory Springer 9781846281815.

Lehmann J, Libchaber A (2008) Degeneracy of the genetic code and stability of the base pair at the second position of the anticodon RNA 14:1264–1269. https://doi.org/10.1261/rna.1029808.

Rietman EA, Karp RL, Tuszynski JA (2011) Review and application of group theory to molecular systems biology Theoretical Biology and Medical Modelling 8:21. https://doi.org/10.1186/1742-4682-8-21.

Sanchez R, Morgado E, Grau R (2005) Gene algebra from a genetic code algebraic structure Journal of Mathematical Biology 51:431–457. https://doi.org/10.1007/s00285-005-0332-8.

Sanchez R, Grau R, Morgado E (2006) A novel Lie algebra of the genetic code over the Galois field of four DNA bases Mathematical Biosciences 202:156–174. https://doi.org/10.1016/j.mbs.2006.03.017.

Ye S, Lehmann J (2022) Genetic code degeneracy is established by the decoding center of the ribosome Nucleic Acids Research 50:4113–4126. https://doi.org/10.1093/nar/gkac171.

